# Are dumbbell stomata unique? Diversified developmental trajectories in sedges and grasses result in partially convergent stomata

**DOI:** 10.64898/2026.05.18.726033

**Authors:** Emilio Petrone-Mendoza, Eleonore Cinti, Maria Rosaria Barone Lumaga, Lara Reale, Salvatore Cozzolino

**Affiliations:** Department of Biology, University of Naples Federico II, Naples 80126, Italy; Department of Agricultural, Food and Environmental Sciences, University of Perugia, Perugia, Italy

**Keywords:** dumbbell-stomata, kidney-stomata, Cyperaceae, Poaceae, developmental-trajectories, cellulose-microfibrils, nuclei, plasmodesmata

## Abstract

Dumbbell-shaped stomata in grasses represent an evolutionary novelty, as their distinctive guard cell morphology is absent from most plant lineages. Stomata exhibit two major morphological forms: the kidney-shaped type found in most plants, and the dumbbell-shaped type that evolved in grasses. Dumbbell-like forms occur in sedges (Cyperaceae), providing an opportunity to examine how changes in developmental trajectories contribute to morphological evolution. By integrating analyses of cellulose microfibril organization, guard cell length to width ratio, and nuclear morphology, we demonstrate partial convergence between sedge and grass stomatal development. Specifically, cellulose microfibril organization in sedges represents an intermediate developmental state between kidney-shaped stomata and the grass dumbbell-shaped stomata. We further document differences in nuclear architecture: in contrast to kidney-shaped stomata, which have rounded nuclei in central guard cell regions, sedge nuclei are partially elongated and localize within bulbous regions, whereas grass nuclei exhibit fully elongated shapes along the cell axis. Notably, we identified secondary plasmodesmata between guard cells in one sedge species, suggesting a convergent route to symplastic communication achieved through secondary plasmodesmata formation rather than the incomplete cytokinesis characteristic of grasses. Together, these findings reveal convergent developmental solutions underlying similar stomatal morphologies.

## INTRODUCTION

The evolution and development of traits associated with differential diversification rates among lineages is an active topic in evolutionary biology. Traits that increase net diversification rates by enabling lineages to exploit new adaptive zones have been termed key innovations (Hodges & Arnold, 1995; Cacho *et al*., 2010; Miller *et al*., 2023; De-Kayne *et al*., 2025). In addition to testing whether a trait confers a higher diversification rate, characterizing the developmental basis helps in explaining the novelty and how it is linked to a new adaptive zone (Moczek, 2008). In particular, comparative data from closely related lineages are essential for identifying the origin and nature of evolutionary novelties. Dumbbell-shaped stomata in grasses provide a compelling model system: their distinctive morphology is not observed in most other lineages (Fig 1), and without comparative data, the evolutionary processes underlying this trait remain unresolved.

**Fig 1.**
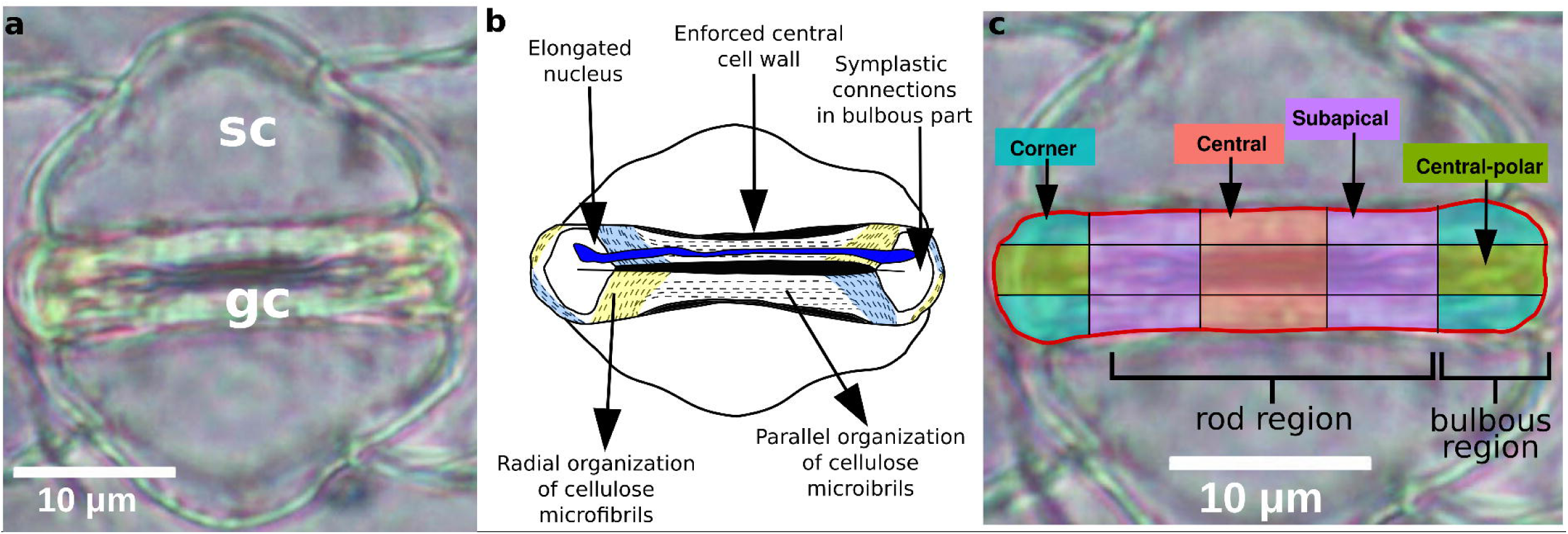
Example of the dumbbell stomata in the grass Zea mays. (A) Light micrograph of a Zea mays stomatal complex. A pair of guard cells (gc) flanked by two subsidiary cells (sc). (B) Schematic representation of the grass stomatal complex and dumbbell-shaped guard cells, highlighting key structural traits (modified from Spiegelhalder & Raissig, 2021). (C) Spatial regionalization of a horizontally oriented pair of guard cells based on a 5×3 grid. The central column corresponds to the central region; the second and fourth columns define the subapical regions; the second row of the first and fifth columns corresponds to the central-polar regions; and the first and third rows of the first and fifth columns define the corner regions. In addition, according to the traditional morphological classification, the central region corresponds to the central rod, the subapical regions encompass the transition between the central rod and the bulbous ends, and the corner regions correspond to the bulbous regions. The stomatal complex represented in A and C is the same.

Dumbbell-shaped stomata develop through a two-stage process. First, the guard mother cells divide to produce a pair of guard cells (GCs) that initially resemble the kidney-shaped stomata typical of most angiosperms. Second, these GC), together with the adjacent subsidiary cells, undergo pronounced elongation (Cleary & Hardham, 1989; Galatis & Apostolakos, 2004; Nunes *et al*., 2020). Concomitant with cell elongation, cellulose microfibrils in the central region of the GCs reorganize from a radial arrangement to one oriented parallel to the pore (Galatis & Apostolakos, 2004), and the nucleus also becomes elongated – sometimes described as dumbbell-shaped (Sack, 1993)– (Fig 1b). GC elongation, microfibril reorganization, and nuclear elongation all occur after GC division and are not observed in kidney-shaped stomata. The observation of extra developmental steps in the grass stomata ontogeny is consistent with a peramorphic trajectory with respect to kidney-shaped stomata, that means the ontogeny of the grass stomata recapitulates the trajectory of the species with the kidney ontogeny adding novel steps to the developmental trajectory. In addition, mature GCs in kidney-shaped stomata typically lack symplastic connections, exhibiting hydraulic independence (Voss *et al*., 2018; Cui *et al*., 2023; Brodersen *et al*., 2025). In contrast, grass GCs maintain symplastic connectivity (Brown & Johnson, 1962; Galatis, 1980; Spiegelhalder & Raissig, 2021; Wilson *et al*., 2025) through incomplete cytokinesis after guard mother cells divide to form paired GCs (Kaufman *et al*., 1970). In addition to grasses, dumbbell-shaped GCs have been described in other families of the order Poales (Tomlinson, 1974; Mishkind *et al*., 1981; Sack, 1993; Rudall *et al*., 2017).

Stomatal formation across Poales is presumed to follow a shared developmental framework involving a basipetal differentiation of the stomatal complexes along the organized cell files in the leaf, beginning with the guard mother cells formation and paracytic subsidiary cell induction (Stebbins & Khush, 1961; Mishkind *et al*., 1981; Croxdale, 2000; Rudall *et al*., 2017). After the subsidiary cells differentiate, the guard mother cells divide and the pore forms between the two GCs. Nevertheless, the morphological and evolutionary diversity of dumbbell-like stomata remains poorly understood. In *Flagellaria indica* (Flagellariaceae), a clade sister to Poaceae, GCs elongate in a dumbbell-like fashion but to a lesser extent than in grasses, and their nuclei remain roundish (Sack, 1993). In the sedges (Cyperaceae) and rushes (Juncaceae), GC elongation is variable, and cellulose microfibrils are reported as predominantly radially oriented (Mishkind *et al*., 1981). Clarifying the evolution and structural characteristics of dumbbell-like stomata is crucial for determining whether, and why, fully developed dumbbell stomata evolved only in grasses.

Sedges in particular provide an ideal system to study the evolution and development of dumbbell-shaped stomata. Sedges are the third largest monocot group, with lineages that have diversified in both tropical (e.g. *Cyperus* spp.) and temperate regions (e.g. *Carex* spp.) (Martín-Bravo *et al*., 2019; Larridon *et al*., 2021; Márquez-Corro *et al*., 2021). Grasses and sedges have also diversified under similar ecological conditions, with elevated speciation rates associated with open, sunny habitats (Elliott *et al*., 2024). Notably, both grasses and sedges also include lineages that independently evolved C4 photosynthesis multiple times (Besnard *et al*., 2009; Sage *et al*., 2011). Furthermore, in some ecosystems, such as African savannas, both grasses *(e.g., Andropogon sp.*) and sedges (*Bulbostylis igneotonsa*) can be dominant components of the vegetation (Medwecka-Kornaś & Kornaś, 1985). Together, the phylogenetic and ecological overlap make sedges a powerful comparative system for testing whether dumbbell-shaped stomata represent a grass-specific innovation or a broader evolutionary convergence within Poales.

In the present work, we compare stomatal morphology across species representing six of the 15 major clades of the Cyperaceae, including members of the Mapanioideae and the Cyperoideae subfamilies (Larridon *et al*., 2021; Cyperaceae Working Group, 2025). We examine the length and width of the GCs, the cellulose microfibril orientation, GC nuclei shape and position (Fig 1). We test the hypothesis that the cellulose microfibril reorganization observed in grasses is associated with post-pore elongation of the GCs. Given that GC elongation in sedges is variable but consistently less pronounced than in grasses, we predict only partial cellulose microfibril reorganization in sedges. We further predict that nuclear shape and position will vary among sedge species and that greater GC elongation will be associated with increased nuclear elongation or with migration into the bulbous region of GCs. Additionally, we report the presence of symplastic connections in the GC bulbous region of one sedge species.

## MATERIALS AND METHODS

### Plant materials

To uncover convergence in the stomata morphology between sedge dumbbell-like stomata and grass dumbbell-shaped stomata, we analyzed 28 species: one eudicot kidney-shaped species, one monocot non-Poales species, one Bromeliaceae, one Typhaceae, and one Juncaceae species, six grass species and 17 sedge species (Table S1). From the Cyperaceae family, we selected species from six different genera including one species from the Mapanoideae subfamily (Table S1, Larridon *et al*., 2021). Plant material came from different sources, such as botanic gardens, or seeds acquired from commercial stores. Our sampling included species from wet tropical forests such as *Didymiandrum stellatum* or *Scirpodendron ghaeri,* to species from seasonally dry tropical forests to species from open and closed temperate climates such as *Carex flava.* Some herbarium specimens were analyzed, though observations were restricted to cellulose microfibril organization due to limited material availability (Table S1).

### Cellulose microfibril organization

To examine cellulose microfibril organization, we employed standard polarized light microscopy. For living material, epidermal peels from the abaxial side of 11 species of the mid-section of fully expanded leaves or culms (hollow stems in sedge and grasses) were mounted in distilled water and observed under a Zeiss Axiolab light microscope (Zeiss, Jena, Germany) equipped with a retardation plate inserted at 45° to a compensator Lamba polarization plate (rotatable +/- 5°). For herbarium specimens, a leaf section was hydrated for at least 10 minutes in distilled water followed by an epidermal peel procedure. Epidermal peels were oriented parallel to the polarization plate. When the crystalline cellulose microfibrils are oriented parallel or perpendicular to the major axis of the retardation plate (45° or 135°), a colour emission is generated, while when the cellulose microfibrils are oriented parallel to the polarizer no color emission is generated, allowing to infer the orientation of the cellulose microfibrils in the stomata. When fresh leaf material was available, we sampled leaves from three individuals and captured images of at least 10 distinct stomata per sample. Due to material limitations in herbarium specimens, only one or two stomata from a single sample were photographed (Table S1). Additionally, primordial leaves (less than 1cm in length) from one *Cyperus alternifolius* individual were sampled to observe cellulose microfibril organization in non-mature GCs. Stomata were photographed with a Nikon Digital Sight DS-L1 (Nikon, Tokyo, Japan).

### Nuclear and organelle organization

To assess nuclear morphology in GCs, we used the fluorescent dye DAPI. Fresh epidermal peels from eight species were obtained with a razor blade and fine forceps and fixed in FAA (50% (v/v) ethanol (95%), 5% (v/v) glacial acetic acid, 10% (v/v) formaldehyde (37–40%), and 35% (v/v) distilled water) for at least 48 h. Samples were rinsed twice in 95% ethanol and placed in 1 μg/ml of Fluoroshield™ DAPI (Sigma-Aldrich), then incubated in the dark for 1 h prior to observation. Images were initially acquired under bright-field illumination and subsequently examined under fluorescence using a DAPI filter, with a light microscope (BX53; Olympus, Tokyo, Japan) equipped with a fluorescence lamp, using a 100× objective. Images were captured using an XC50 camera (Olympus, Tokyo, Japan).

To examine the presence of symplastic connections in the bulbous region of the GCs we prepared samples from *Zea mays*, *Cyperus esculentus,* and *Cyperus alternifolius* to be observed using transmission electron microscopy (TEM). These species were selected based on differences in guard cell organization and nuclear positioning. In *C. alternifolius*, DAPI staining consistently revealed nuclei predominantly localized in the bulbous regions of the guard cells; this species was therefore selected to investigate a potential relationship between nuclear positioning and symplastic connectivity. In contrast, *C. esculentus* exhibited variable nuclear morphology and positioning, with nuclei observed either in the central region or in the bulbous regions of the guard cells, providing an intermediate reference for assessing the association between nuclear position and symplastic connectivity. Small cross-sections of leaves were fixed in 2.5% (v/v) glutaraldehyde in 0.1 M phosphate buffer at pH 7.2 and kept at 4° C overnight. Then the samples were post-fixed in 1% (w/v) osmium tetroxide (OsO_4_) and 0.8% KFeCN in phosphate buffer at room temperature for 1.5 h followed by an alcohol dehydration series to finally embed the samples in Spurr’s resin. Sections were cut at 70 nm using a Reichert-jung supernova ultramicrotome. Ultrathin sections were collected on a 200-mesh uncoated copper grids, stained in Uranyl Acetate Replacement UAR (Electron Microscopy Science) and post-stained in lead citrate (CeSMA, section of Microscopy, University of Naples Federico II). Sample examination was done on a Tecnai G2 S-Twin TEM. We only observed two stomata from one sample per species with TEM microscopy.

### Image and statistical analysis

#### Stomata size measurements, cellulose microfibril distribution, and nuclei distribution

To test whether guard cell (GC) elongation –quantified as the length to width ratio– differed among kidney-shaped stomata, sedge dumbbell-like stomata, and grass dumbbell-shaped stomata, we performed a one-way analysis of variance (ANOVA). When the ANOVA indicated significant differences, we conducted Tukey’s honestly significant difference (HSD) post hoc tests to compare all pairwise group means. Compact letter displays summarizing statistically significant differences among groups were generated using the *cld* function from the multcomp package (Piepho, 2004) in the R statistical environment (R Core Team, 2023).

To quantify the spatial distribution of cellulose microfibril orientation within guard cells, processed images were converted to grayscale and segmented using Otsu’s thresholding method applied within each stomatal ROI. This segmentation isolated birefringent regions corresponding to cellulose microfibrils oriented either parallel (45°) or perpendicular (135°) to the retardation plate. Foreground pixels in the resulting binary images were used as a proxy for cellulose microfibril birefringence. Each stomatal mask was subdivided into a regular grid consisting of five columns and three rows of equal size, yielding 15 subregions aligned with the major axis of the stomatal aperture (Figs 1C, 2 and 3). These subregions were grouped into four functional categories: (i) the central region (third column), (ii) subapical regions (second and fourth columns), (iii) central-polar regions (second row of the first and fifth columns), and (iv) corner regions (first and third rows of the first and fifth columns). For each category, the total birefringent area was quantified as the proportion of foreground (binarized) pixels relative to the total area of the corresponding subregion. We performed the analysis using OpenCV and Scikit-image libraries from Python (Bradski, 2000; Van Der Walt *et al*., 2014).

**Fig 2.**
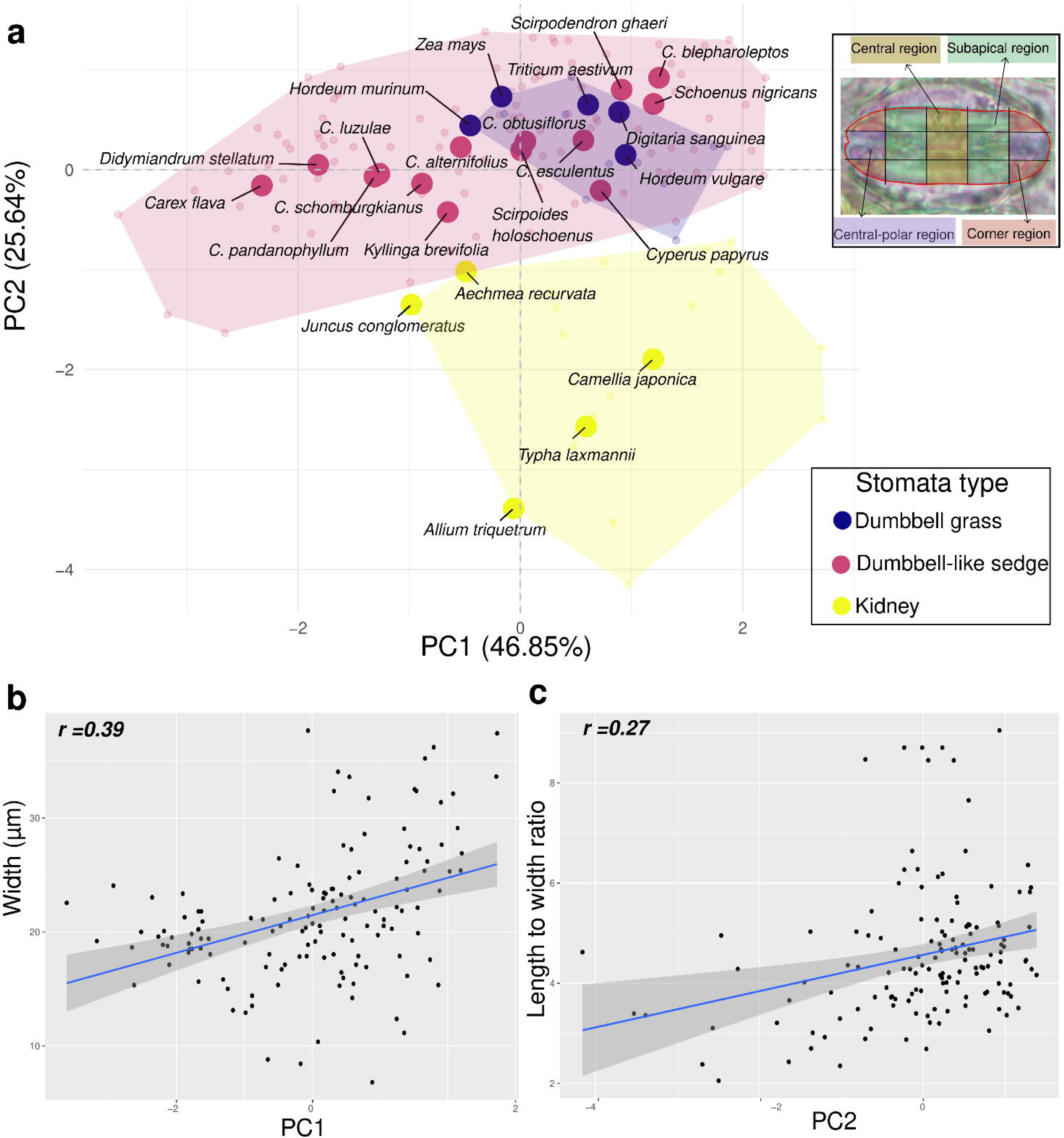
The GC length to width ratio is related to the reorganization of cellulose microfibrils in stomata of grasses and sedges. (A) PCA of the amount of birefringent signal per substomatal region. Inset of the Fig showing the different subregions defined per stomata to quantify distribution of birefringent light per region as a proxy for cellulose microfibril organization. Large circles represent the mean value per species. (B) Correlation observed between stomata width and the PC1 (*p < 0.00001* ) and (C) correlation between length to width ratio and the PC2 (*p = 0.008*).

To identify patterns of cellulose microfibril distribution across species, we performed a PCA on the relative birefringent area measured in each of the four stomatal regions (central, subapical, central-polar, and corner). Prior to PCA, all variables were scaled to unit variance. Because the transition from radial to parallel cellulose microfibril orientation in grasses is associated with guard cell elongation, we examined correlations between the first two principal components and stomatal length, width, and length to width ratio. We tested whether the principal components reflecting reduced birefringence in central regions and increased birefringence in polar and corner regions were associated with more elongated guard cells.

To quantify nucleus shape and position within stomata, images were initially converted to binary using Otsu’s thresholding method. In a subset of images, elevated background fluorescence prevented reliable automated segmentation of the nucleus. In these cases, nuclei were manually segmented using the polygon selection tool in ImageJ, and background signal was removed using the Clear Outside function. The resulting manually curated binary masks were used for all subsequent analyses. Nuclear position was quantified using the same grid-based regional classification applied in the cellulose microfibril analysis, allowing direct comparison of nuclear localization patterns across stomatal regions.

To test whether nuclear distribution differed among regions and species, we fitted a beta regression model using the R package *glmmTMB* (Brooks *et al*., 2017). The relative nuclear area exhibited a non-normal distribution and was strongly biased toward zero; therefore, a beta regression with a logit link was used to model proportional data bounded between 0 and 1. Species, region, and their interaction were included as fixed effects. A zero-inflated component was incorporated to account for excess zeros. Model selection was based on likelihood ratio tests, which indicated that the model including the species × region interaction provided a significantly better fit than the additive model. Estimated marginal means were therefore calculated from the interaction model using the *emmeans* package in R (Lenth *et al*., 2021). Pairwise comparisons among regions within species were conducted using compact letter displays implemented with the *cld* function, with p-values adjusted using the Tukey method to account for multiple comparisons (Piepho, 2004). Confidence intervals were back-transformed from the logit scale, and all statistical tests were performed on the log-odds ratio scale. Finally, to identify patterns of nucleus location and shape, and GC shape, we performed a PCA on the relative area of nucleus across the four stomatal regions (central, subapical, central-polar, and corner), nucleus area, length and width, and the GC length to width ratio. Prior to PCA, all variables were scaled to unit variance.

#### Code and data availability

Custom scripts for image processing and analysis were developed by the authors, with assistance from the large language model ChatGPT (OpenAI, 2025) (version GPT-5 mini) for code structuring and debugging. OpenCV and Scikit-image libraries from Python were used (Bradski, 2000; Van Der Walt *et al*., 2014). All analyses, validation steps, and biological interpretations were performed by us. The code used for image processing developed in Python (Van Rossum & Drake, 2009) and statistical analyses developed in R (R Core Team, 2023 will be made publicly available on GitHub upon acceptance. All images analyzed in this study will be deposited in a publicly accessible repository (Link available once accepted).

## RESULTS

A one-way analysis of variance (ANOVA) detected a significant effect of stomatal type on guard cell elongation (length to width ratio; F_2,360_ = 105.1, *p* < 0.001). Post hoc comparisons indicated that kidney-shaped, sedge dumbbell-like, and grass dumbbell-shaped stomata differed significantly. Kidney-shaped stomata had length to width ratios of 1.53 (± 0.21), sedge dumbbell-like stomata had values of 2.88 (± 0.11), while grass dumbbell-shaped stomata had values of 3.25 (± 0.29) (Fig S1).

### Cellulose microfibril organization in sedge stomata shows an intermediate pattern with respect to grass dumbbell stomata

Cellulose microfibril orientation in stomata of sedges was variable, ranging from radially oriented around the pore –typical of kidney-shaped stomata– to parallel oriented along the axis in the central region, as seen in dumbbell-shaped stomata (Fig S2). The first component of the PCA explained 46.85% of the variation while the second component explained 25.64% of the total variation of birefringence signal across stomatal regions. The first component captured a positive correlation between percentage of birefringence in the central and subapical regions, a positive correlation between birefringence in the corner and central-polar regions and a negative correlation between birefringence in the central–subapical pair and the corner–central-polar pair (Fig 2a). The second component reflected the overall level of birefringence across regions, distinguishing stomata with high birefringence in all regions from those with generally low birefringence. Species having kidney-shaped stomata tended to have birefringence in all regions, while grasses had birefringence in the central-polar and corner regions (Fig 2a). Within sedges, species such as *Carex flava*, had birefringence in the subapical regions and no birefringence in the central-polar and corner regions (Fig 2a, Fig S2). Species such as *Cyperus esculentus* and *Cyperus alternifolius* had both stomata with the same pattern as *Carex flava* and stomata with birefringence in the central-polar and corner regions without birefringence in the central region as in other grasses (Fig 2a, Fig S2). We found a positive correlation between PC1 and stomatal width (r^2^= 0.39, *p-value <*0.001, Table S2, Fig 2b) and PC2 and the length to width ratio (r^2^= 0.27, *p-value =* 0.008). Consistent with cellulose microfibril orientations reported for immature grass stomata (Galatis & Apostolakos, 2004), we observed low birefringence levels or kidney-like birefringence patterns in immature *C. alternifolius* stomata. (Fig S3).

### Nuclear shape and organelle organization in the sedge bulbous stomata region

The DAPI staining revealed that the nuclear position in the GCs of sedges varies between species (Fig 3, Fig S4). In the PCA, the first component explained 37.95% of the variation, and the second component explained 26.48% of the variation. High PC1 values corresponded to larger and more elongated nuclei, especially located in central and corner subregions, while low PC1 values corresponded to smaller nuclei, often associated with subapical regions (Fig 3a). The second component captured, mainly, variation in the localization of the nuclei within GCs and the length to width-ratio. Nuclei localization in central, subapical regions, was negatively correlated with corner and central-polar localization and length to width ratio. Therefore, stomata with higher length to width ratios tended to have nuclei located in the corner and central-polar regions. Sedge species tended to have smaller nuclei with variation in nuclei location. Quantification of relative nuclear areas across regions revealed species-specific spatial patterns. In *Cyperus glaber, Carex flacca, Didymiandrum stellatum*, and *Camellia japonica,* nuclei were predominantly localized in the central and subapical regions. In contrast, in *Cyperus esculentus, Cyperus alternifolius*, and *Carex sylvatica*, relative nuclear area was evenly distributed across all subregions. In *Lolium perenne*, nuclei were more abundant in the central–polar regions (Fig 3b, Fig S2).

**Fig 3.**
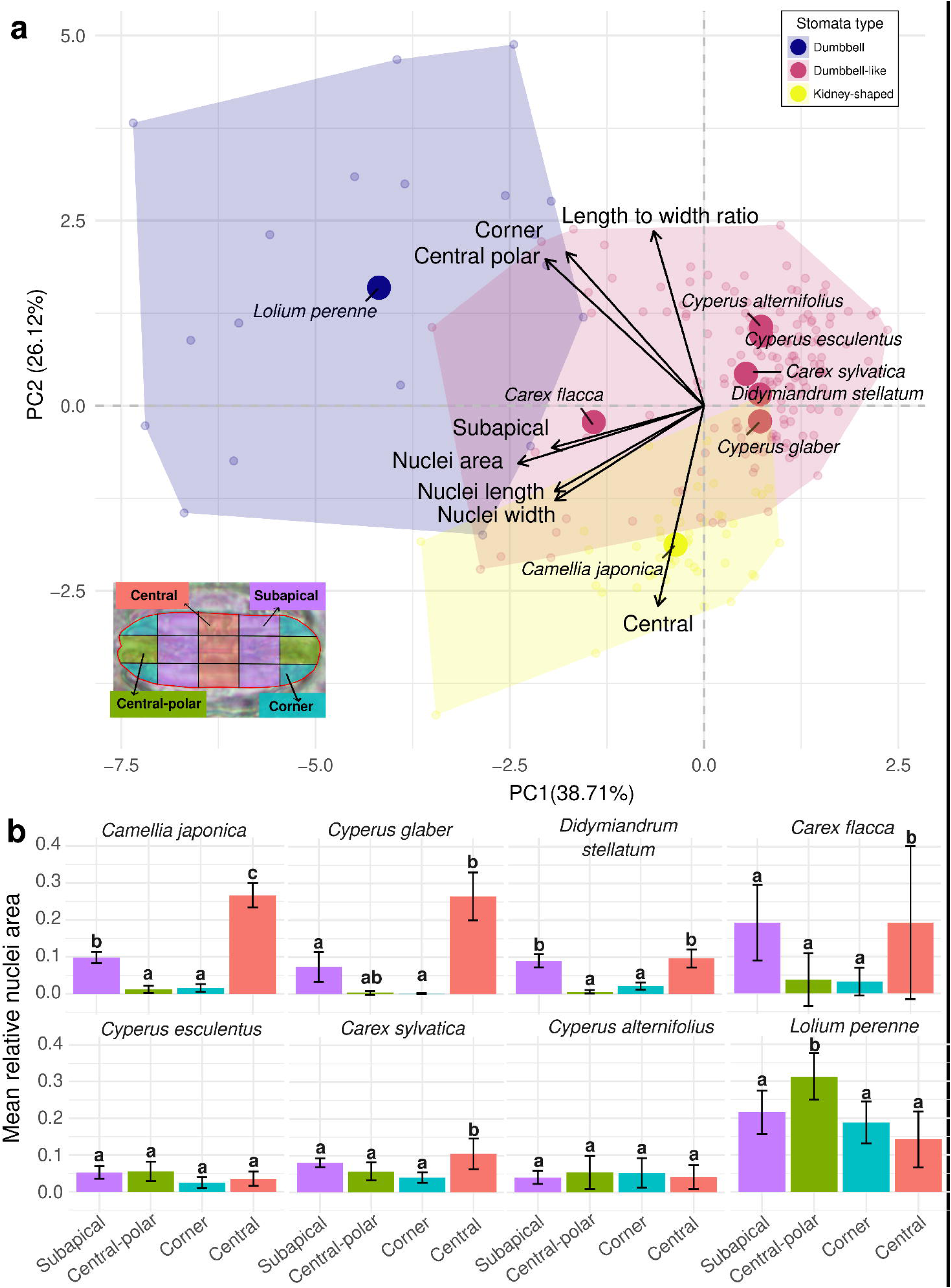
Nuclei shape variation across sedge species. (A) Principal component analysis (PCA) of nuclei traits measured across stomatal subregions. Small points represent individual observations, colored by stomatal morphotype (kidney-shaped, sedge-like, grass dumbbell), while larger points represent the mean values by species. Convex hulls indicate the multivariate space occupied by each morphotype. Vectors show loadings of nuclei traits (length, width, area) and spatial subregions (central, central-polar, corner, subapical). PC1 primarily reflects variation in nuclei size and elongation, separating grass dumbbell stomata from kidney-shaped and sedge stomata. PC2 captures differences associated with polar versus central nuclei organization. Together, the PCA reveals distinct multivariate nuclei architectures among stomatal morphotypes and along the stomatal axis. (B) Barplot showing the mean relative area of nuclei by region. Different letters indicate significant differences among regions within each species based on Tukey-adjusted pairwise comparisons of estimated marginal means.

As reported in other studies (Louguet *et al*., 1990; Galatis & Apostolakos, 2004; Wilson *et al*., 2025), we observed symplastic continuity between GCs in *Zea mays* (Fig 4). On the contrary, *C. alternifolius* and *C. esculentus* did not show incomplete cytokinesis (Fig 5, and 6), but we observed cell wall thinning in the bulbous region of *C. alternifolius* (Figs 5). Concomitant with the cell wall thinning, a pit field of secondary plasmodesmata was observed (Figs 5b). In addition, transmission electron microscopy images of *C. alternifolius* revealed that the cellulose microfibrils in the central part of the GCs, towards the bulbous region, are oriented at 45° near the ventral region, while a more parallel orientation appears toward the dorsal region resembling the grass dumbbell orientation (Figs 5c). At the interface between the bulbous and central regions, a radial orientation was also evident. In contrast to *C. alternifolius*, we did not observe plasmodesmata in the stomata of *C. esculentus,* instead we observed a thick cell wall between the GCs (Fig 6). We can also distinguish a mixture of endoplasmic reticulum vesicles and vacuoles (v) and part of the nucleus of *C. esculentus*.

**Fig 4.**
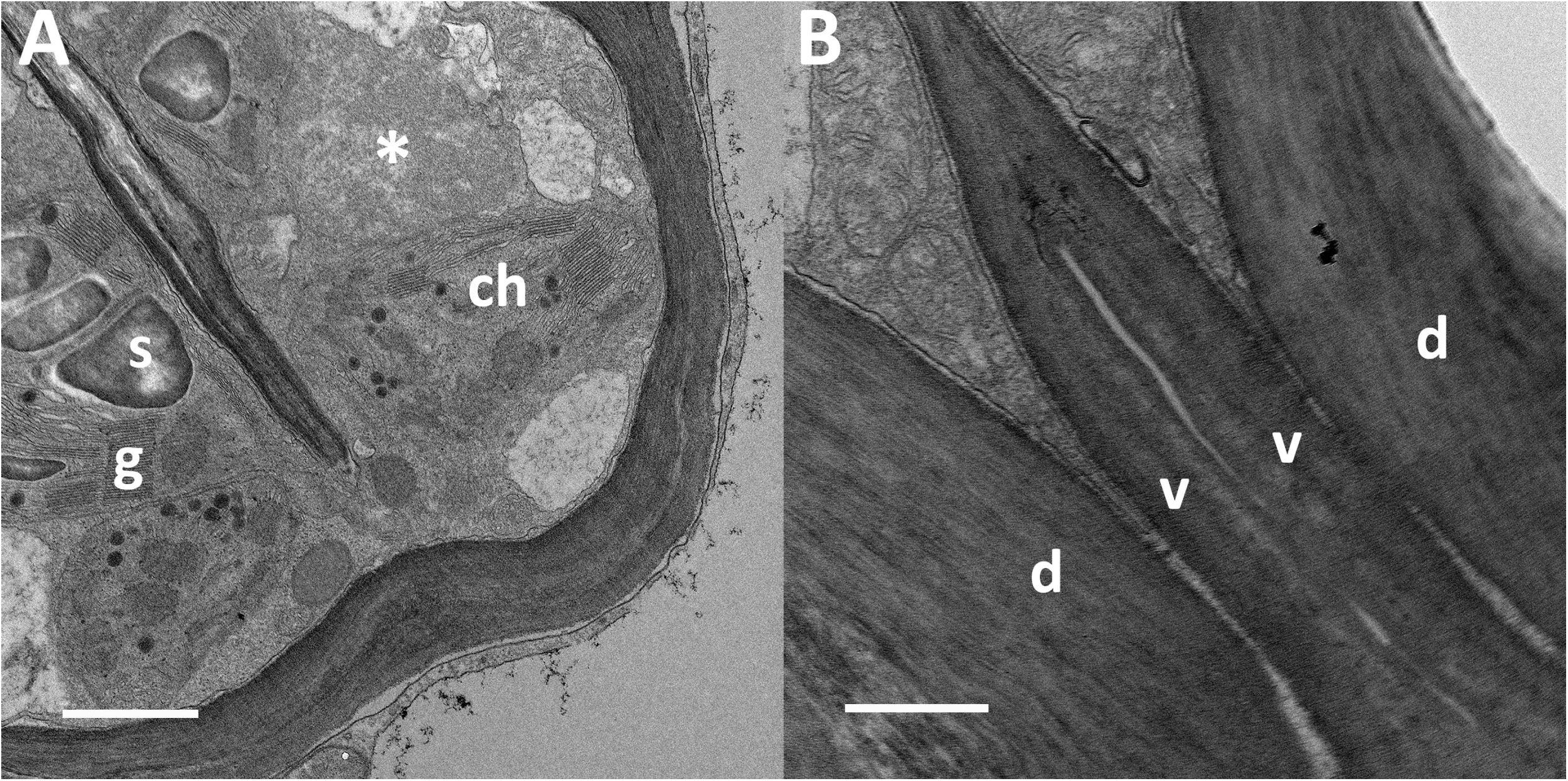
Transmission electron microscopy micrographs of *Zea mays* showing the ultrastructure of the dumbbell stomata. (A) Bulbous region showing large chloroplasts (ch) with grana (g) and starch grains (s). The asterisk highlights part of the nucleus. (B) Thick cell wall engrossment in the area delimiting the bulbous and central region of the GCs. Note the parallel orientation of the cellulose fibrils in the cell wall both dorsal (d) and ventral (v) of the central region of the GCs. Scale bars: A 1µm, B 500 nm.

**Fig 5.**
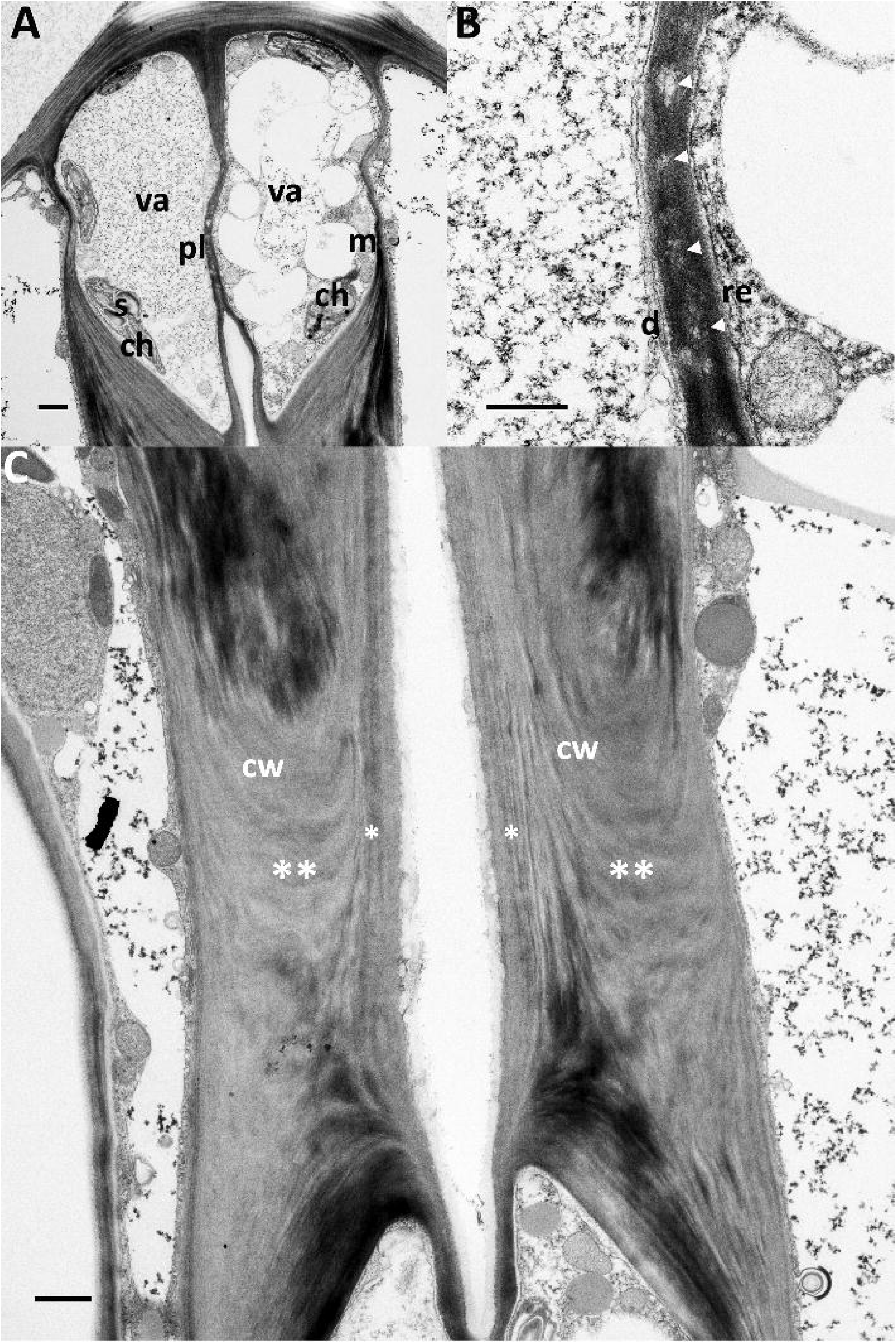
Transmission electron microscopy micrographs of *Cyperus alternifolius* showing the ultrastructure of GCs. (A) Bulbous region showing vacuole(va), mitochondria (m), and chloroplasts (ch) containing small starch granules (s), and plasmodesmata (pl) between the cell wall of the GCs. (B) Close up to the cell wall between the two GCs, endoplasmic reticulum (er) and desmotubules (d) associated with plasmodesmata (arrowheads) can be observed. (C) Cellulose fibrils in the cell wall (cw) show a radial organization (**) in the central part of the GCs. The ventral cell wall shows a parallel orientation (*) of the cellulose fibrils. Scale bars: A 1µm, B 500 nm, C 1µm.

**Fig 6.**
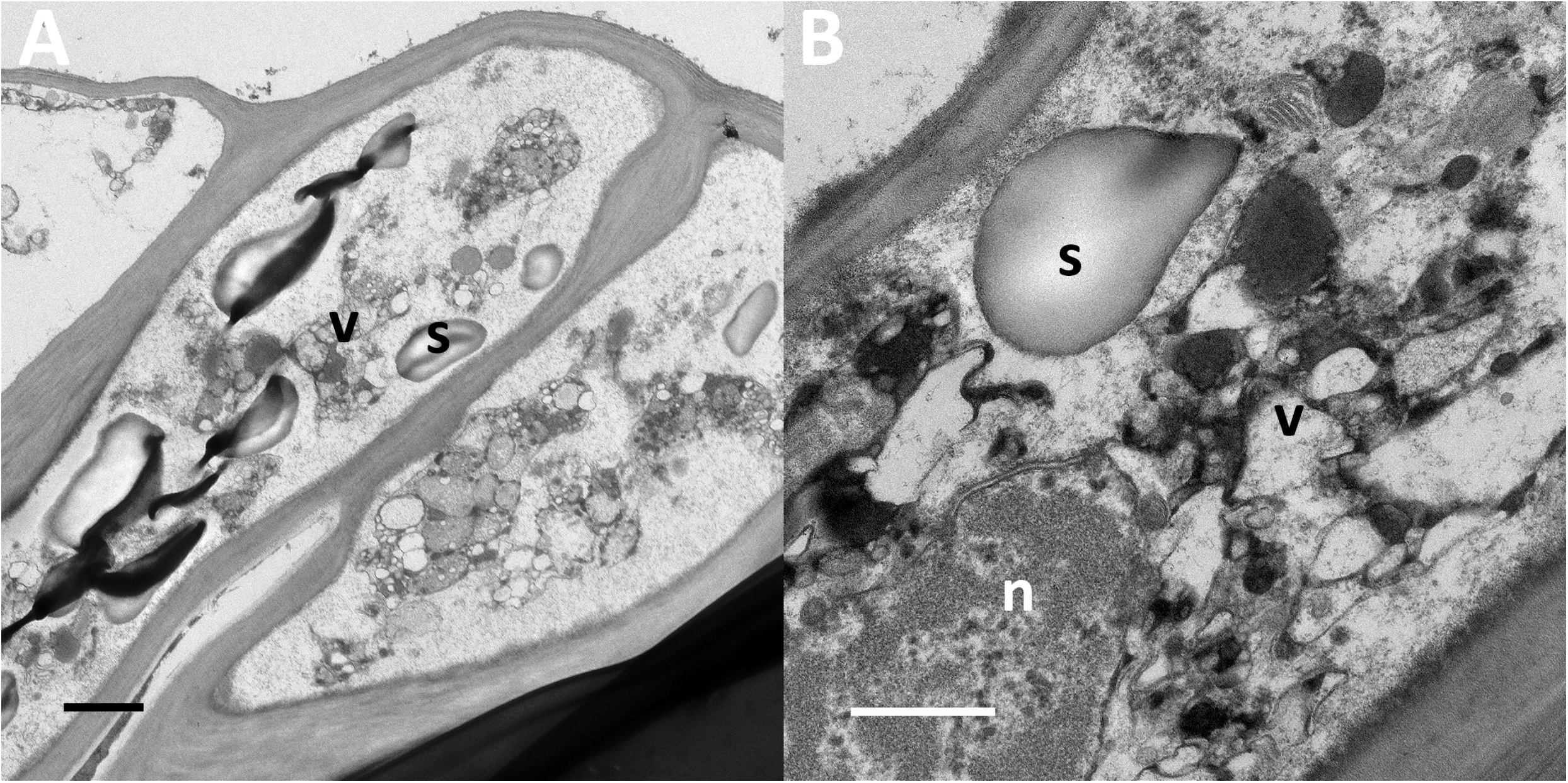
Transmission electron microscopy micrographs of *Cyperus esculentus* showing the ultrastructure of the GCs. (A) Bulbous region showing starch granules (s), and dilated endoplasmic reticulum vesicles (v). No symplastic continuity is evident between the two GCs. Slight thinning of the cell wall is present. (B) Central portion of the GC showing starch granules (s), endoplasmic reticulum vesicles or vacuoles (v) and part of the nucleus (n). Scale bars: A 1µm, B 500 nm.

Although DAPI staining cannot directly confirm the presence of secondary plasmodesmata, the nuclei of paired guard cells were consistently observed in close contact in the bulbous region of *C. sylvatica, C. alternifolius* (Fig S5), and in the grass *L. perenne* (Fig S6), suggesting potential symplastic connectivity. In contrast, such nuclear proximity was not observed in the kidney-shaped species *C. japonica* (Fig S7).

## DISCUSSION

In the present work, we show that the GC length to width ratio in sedges has intermediate values between kidney-shaped and grass dumbbell-shaped stomata. We also show that cellulose microfibrils in sedges rearrange partially in the central-rod region of the guard cells, showing an intermediate phenotype with respect to grasses. We also show that nuclear localization in sedges tends to be located in the bulbous region of the cells. The position of the nucleus within the GCs could be linked to GC elongation. In addition, we report the presence of secondary plasmodesmata between guard cells of *Cyperus alternifolius*. Together, our analyses show partial convergence of stomatal traits between sedges and grasses.

### Shared early stomatal developmental trajectory between sedges and grasses

The succession of organismal features from early stages to maturity and beyond are known as developmental trajectories (Olson, 2024). Moreover, when developmental trajectories are mapped across species, changes in the timing of developmental events—a process known as heterochrony—can be inferred (Alberch *et al*., 1979; Olson, 2020; Onyenedum & Pace, 2021). Within this framework, paedomorphic descendants resemble early stages of the hypothetical ancestral trajectory, whereas peramorphic descendants complete all developmental steps and undergo additional modifications during ontogeny (Alberch *et al*., 1979; Buendía-Monreal & Gillmor, 2018; Petrone-Mendoza *et al*., 2023).

The ancestral stomatal type for Poales consists of two GCs and two paracytic subsidiary cells (Rudall *et al*., 2017). In Poales, stomatal complexes develop from epidermal cell files organized in axial files along the leaf (Sajo & Rudall, 1999; Rudall *et al*., 2017; Fogaroli Corrêa *et al*., 2023). In both sedges and grasses, stomatal development begins with a transverse division of an epidermal cell file that produces the guard mother cell; subsequently, a pair of lateral subsidiary mother cells is specified and divides to generate the subsidiary cells. Only after these events does the guard mother cells divide to form the two GCs. Thus, sedges and grasses share an early stomatal developmental trajectory that likely predates their divergence within Poales.

### Divergent and parallel trajectories in GC division and symplastic connectivity

A key developmental difference in stomatal development emerges during GC cytokinesis. In grasses, cytokinesis is incomplete (see Fig 16 in Kaufman *et al*., 1970), leaving a discontinuity in the GC wall and maintaining symplastic continuity that may facilitate turgor equilibration and stomatal movement (Mumm *et al*., 2011; Durney *et al*., 2023; Wilson *et al*., 2025). Although this observation is based on a limited number of species, it suggests that incomplete cytokinesis is a novelty in the Poales stomata development. Available evidence in sedges, indicates complete cytokinesis after guard mother cell division, pointing to a divergence in stomatal development from grasses.

Following GC formation, grasses undergo a pronounced elongation accompanied by central wall thickening, nuclear elongation, and reorganization of cellulose microfibrils into a pore-parallel configuration (Galatis, 1980; Galatis & Apostolakos, 2004). Across the taxa examined here, guard cell elongation varied among stomatal morphotypes (kidney-shaped, sedge, and dumbbell-shaped stomata; Fig S1). Species from Typhaceae and Juncaeae, although sparsely sampled, displayed comparatively low elongation and retained a microfibril organization resembling that of kidney-shaped stomata. In contrast, sedges occupied an intermediate range, with length to width ratios greater than those of kidney-shaped stomata but lower than those of grasses. In some cases, this intermediate morphology resembles early developmental stages described in grasses. For example, *Carex flava* showed limited GC elongation and only partial radial cellulose microfibril patterns with high subapical birefringence, whereas some stomata of *C. esculentus* show reduced birefringence in the central region, consistent with more pronounced microfibril reorganization.

These observations are consistent with the possibility that variation in stomatal morphology across these lineages reflects differences in the timing or extent of shared developmental processes. The observed correlation between GC elongation (length to width ratio) and microfibril alignment (PC2) (Fig 3) supports the existence of coordinated developmental changes, although additional analyses of cell wall composition will be necessary to clarify the mechanistic basis of these patterns. Furthermore, we saw kidney-like microfibril patterns in non-mature stomata of *C. alternifolius* (Fig S3), consistent with the early developmental stages of grasses (Galatis, 1980; Galatis & Apostolakos, 2004).

Under this interpretation, sedges may represent an intermediate condition along a continuum of guard cell elongation and cellulose microfibril reorganization, potentially compatible with heterochronic shifts. However, given the limited taxonomic sampling across Poales, this hypothesis remains tentative and will require broader comparative developmental data to be rigorously tested (Fig 7). Additionally, cell wall composition analyses will be required to interpret the uneven distribution of wall material across the GCs (Shtein *et al*., 2017).

**Fig 7.**
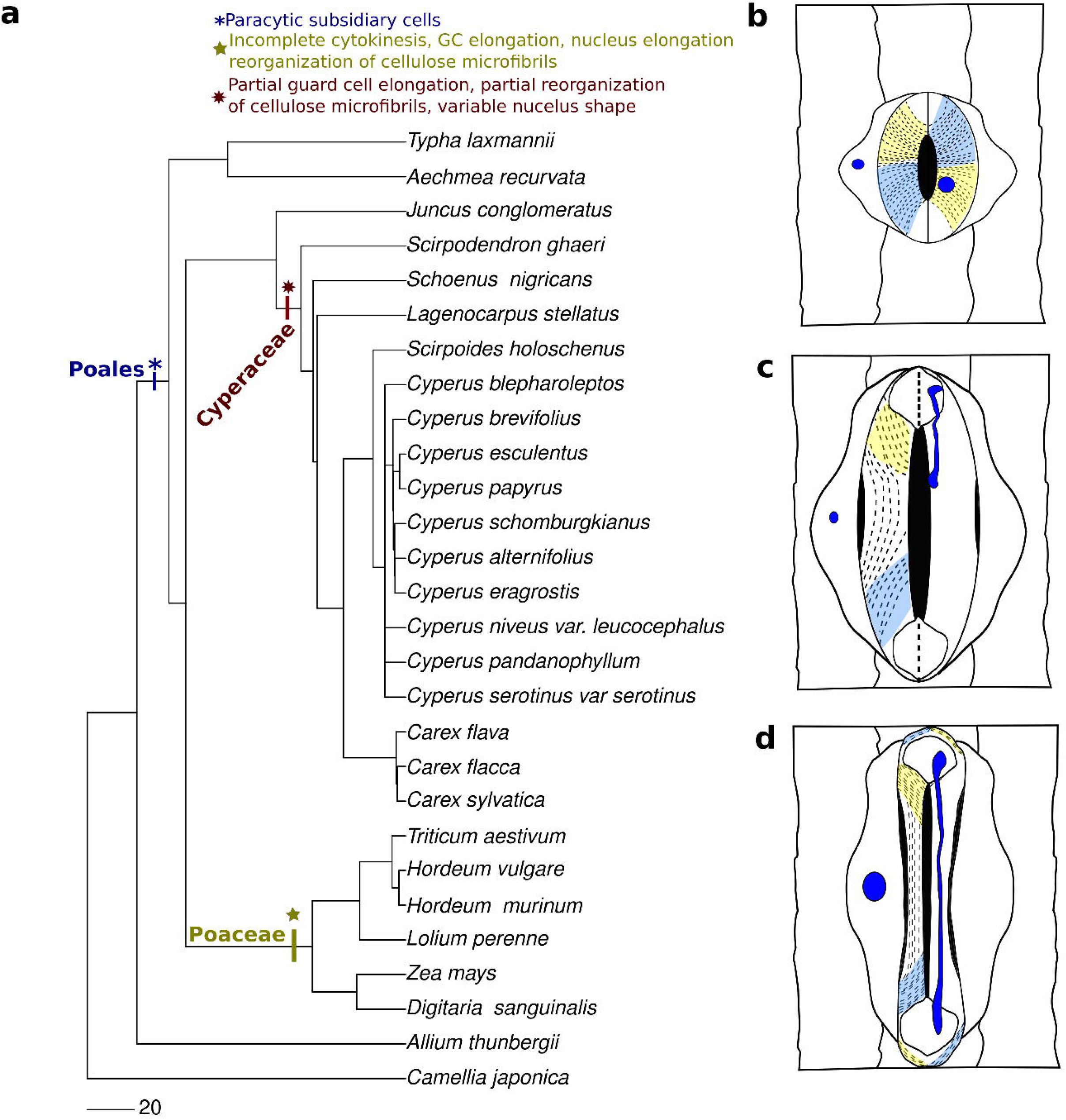
Developmental and cellular traits associated with stomatal diversity across Poales. (**a**) Simplified phylogenetic relationships among Poales families including the sampled taxa, as well as representative lineages with kidney-shaped stomata. This panel is intended to provide phylogenetic context and does not represent a reconstruction of trait evolution. The phylogenetic tree was constructed using the phylomaker R package (Jin & Qian, 2022) (**b**) Traits observed in stomata of Poales species outside Cyperaceae and Poaceae. Paracytic subsidiary cells are present, guard cells (GCs) are kidney-shaped, cellulose microfibrils are radially oriented with respect to the pore, and nuclei are generally spherical and centrally located. (**c**) Traits observed in stomata of Cyperaceae species. Guard cells show moderate elongation relative to other Poales families, cellulose microfibrils are partially rearranged in the central region, and nuclear shape and position are variable, frequently located toward the bulbous regions of the GCs. (**d**) Traits observed in dumbbell-shaped stomata of grasses (Poaceae). Guard cells are highly elongated, cellulose microfibrils are predominantly oriented parallel to the pore, and nuclei are elongated. Colored regions represent domains of cellulose microfibril orientation, and blue circles indicate nuclei.

### Partial parallelism in nuclei shape and position

Nuclear morphology in sedge GCs is variable. In contrast to grasses—where elongated nuclei have been consistently reported (Kaufman *et al*., 1970; Galatis, 1980; Sack, 1993)—sedge nuclei may be rounded, partially elongated, or positioned in different regions of the cell, including the bulbous ends. In species such as *C. alternifolius, C. esculentus,* and *C. sylvatica* nuclei are frequently located in the bulbous region. In grasses, guard cell shape and reduction of cell volume due to cell wall deposition in the rod-central region (Brown & Johnson, 1962) suggesting that cell shape is linked with nuclei elongation. Quantitative analyses of guard cell volume and shape will be required to clarify this relationship.

We also observed plasmodesmata between guard cells in *C. alternifolius,* localized in the bulbous region. In contrast to most seed plant stomata —where plasmodesmata between guard cells are typically lost during development (Willmer & Sexton, 1979; Zhao & Sack, 1999; Cui *et al*., 2023)—these connections occurred in clusters (pit fields), consistent with secondary plasmodesmata formed during cell wall expansion (Tee & Faulkner, 2024). The form of the plasmodesmata observed in Fig 6 suggests that plasmodesmata desmotubules are not straight but branched or funnel-like. However, plasmodesmata were not observed in TEM images of *C. esculentus*, which may reflect developmental stage or species-specific variation. DAPI staining cannot directly confirm the presence of secondary plasmodesmata, but the nuclei of paired guard cells consistently observed in close contact in the bulbous region of *C. sylvatica, C. alternifolius* (Fig S5), and in the grass *L. perenne* (Fig S6), suggesting potential symplastic connectivity further suggesting connections in the bulbous regions of sedge stomata. Given the limited sampling, the distribution and functional significance of these connections remain unclear and will require broader ultrastructural and molecular characterization.

### Morphological parallelisms and its functional implications in dumbbell stomata

Grass dumbbell stomata have been described as exhibiting faster stomatal kinetics in response to fluctuating light conditions and having higher water use efficiency compared with kidney-shaped stomata (McAusland *et al*., 2016; Lawson & Vialet-Chabrand, 2019). Although recent work indicates that individual anatomical traits, including GC type, are only weak predictors of light-induced changes in stomatal conductance, dumbbell stomata nonetheless display faster opening and closing dynamics than kidney-shaped stomata (Woning *et al*., 2025). This enhanced responsiveness has been attributed, at least in part, to the higher ratio of pore area to total guard cell area, which may enable higher potential stomatal conductance (Franks & Farquhar, 2007).

Within grasses, dumbbell-shaped stomata are thought to represent an ancestral condition and are retained across lineages that have repeatedly transitioned into open, high-light environments (Gallaher *et al*., 2022; Elliott *et al*., 2024). In this context, the functional properties of dumbbell stomata may have been co-opted as an exaptation, facilitating rapid stomatal responses and high physiological performance under these environmental conditions. In a comparable ecological context, sedges also include species occupying open habitats (Elliott *et al*., 2024), and some exhibit dumbbell-like stomatal morphologies. If these morphologies arose under similar selective regimes, an open question is whether dumbbell-like stomata in sedges confer physiological advantages comparable to those of grass dumbbell stomata, such as faster stomatal kinetics or enhanced water-use efficiency. Future studies integrating detailed mapping of stomatal morphologies with climatic and ecological variables will be crucial to clarify the selective advantages and physiological performance associated with dumbbell and dumbbell-like stomata.

Moreover, the traits observed in sedges are consistent with the possibility that grass dumbbell-shaped stomata represent a derived developmental condition, associated with specific changes during development, such as incomplete cytokinesis. A study analyzing the evolution of transcription factors involved in stomatal development in sedges identified a potential loss of *SCRM2* (Menezes *et al*., 2025), raising the possibility that *SCRM2*, in conjunction with *FAMA*, contributes to a novel developmental trajectory in grasses, including incomplete cytokinesis, guard cell elongation, cellulose microfibril reorganization, and nuclear elongation. However, the extent to which these genetic differences underlie the observed developmental patterns remains to be tested.

In conclusion, this study suggests that the developmental and cellular basis of dumbbell-shaped stomata in sedges partially overlaps with that described in grasses, while also revealing important lineage-specific differences. Our results are consistent with the hypothesis that variation in stomatal morphology across Poales may arise from modifications of shared developmental pathways, potentially including heterochronic shifts, although broader comparative sampling will be required to evaluate this framework (Fig 7). This work highlights the value of integrating developmental, cellular, and comparative approaches to understand stomatal evolution and provides a foundation for future studies aimed at identifying the genetic and biomechanical drivers (Tan *et al*., 2025) underlying stomatal shape diversity and testing the extent to which convergent pathways have contributed to the evolution of dumbbell-shaped stomata in monocots.

## ABBREVIATIONS

GCs: guard cells

## ACKNOWLEDGMENTS

We would like to acknowledge Clemens Bayer from Palm Garden of Frankfurt and Marc Reynders from Meise Botanic Garden (BR) for providing sedge material to perform part of the analysis. We want to acknowledge Sergio Sorbo, CeSMA, University of Naples Federico II, for his technical assistance in Electron Microscopy techniques. We would also like to thank Mark Olson for the comments on the manuscript. This research was supported by the National Recovery and Resilience Plan (NRRP), Mission 4 Component 2 Investment 1.4 - Call for tender No. 3138 of 16 December 2021, rectified by Decree n.3175 of 18 December 2021 of Italian Ministry of University and Research funded by the European Union – NextGenerationEU; Award Number: Project code CN_00000033, Concession Decree No. 1034 of 17 June 2022 adopted by the Italian Ministry of University and Research, CUP H43C22000530001, Project title “National Biodiversity Future Center - NBFC” and by Italian Ministry of University and Research (MUR) program PRIN 2020 “The biochemical and diffusive optimisation of photosynthesis: evolutionary implications for the development of climate resilient productive plants (EvoPlant)”.

## AUTHOR CONTRIBUTIONS

Conceptualization, EPM and SC; methodology, EPM, EC, and MRB; Investigation, EPM, EC, and MRB; data curation, EPM and EC; formal analysis, EPM and EC; writing—original draft, EPM, EC, and SC; writing—review & editing, EPM, EC, MRB, LR and SC; funding acquisition, SC and ; resources, SC; supervision, SC.

## DATA AVAILABILITY

### Data and code availability

- Light polarization microscopy images and fluorescence microscopy images have been deposited at [data-type-specific repository] as [Database: accession number] and are publicly available as of the date of publication.
- All original code has been deposited at [repository] and is publicly available at [DOI] as of the date of publication.
- Any additional information required to reanalyze the data reported in this paper is available from the lead contact upon request.

## DECLARATION OF GENERATIVE AI AND AI-ASSISTED TECHNOLOGIES

During the preparation of this work the authors used ChatGPT for assistance in code structuring and debugging. After using this tool, the authors reviewed and edited the content as needed and take full responsibility for the content of the published article.

