## Supplementary Material for "Are dumbbell stomata unique? Diversified developmental trajectories in sedges and grasses result in partially convergent stomata"

The following Supporting Information is available for this article:

**Supplementary figures**

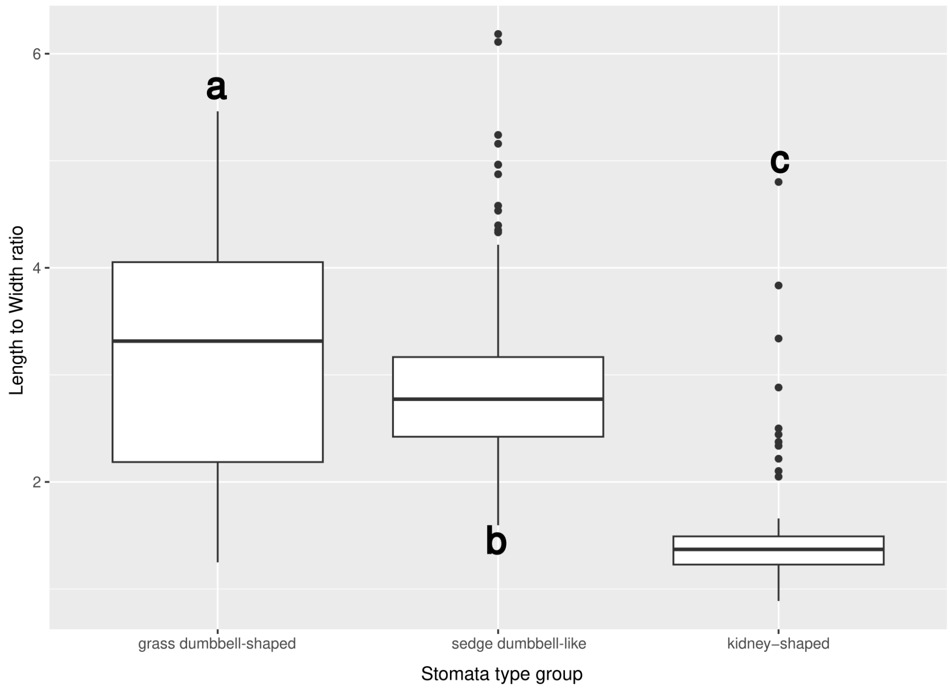


**Fig. S1** Guard cell (GC) elongation, measured as the length-to-width ratio, differed significantly among stomatal morphotypes ($F_{2,360} = 105.1$, $p < 0.0001$). Kidney-shaped stomata showed the lowest elongation (1.53 ± 0.21), whereas sedge stomata (2.88 ± 0.11) and grass dumbbell-shaped stomata (3.25) exhibited substantially higher elongation. Grass dumbbell-shaped stomata were significantly more elongated than sedge stomata. Letters denote statistically significant differences among groups following post-hoc comparisons.

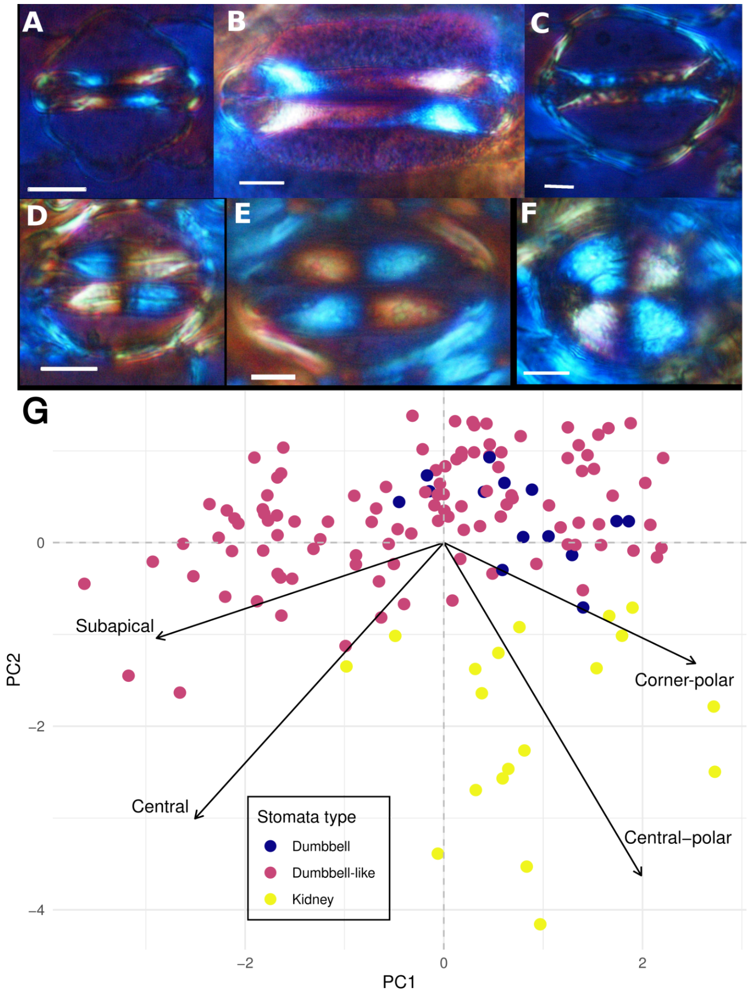


**Fig S2.** Cellulose microfibril organization in sedges show variable patterns between dumbbell stomata and kidney stomata. In the dumbbell stomata of (A) Zea mays and (B) Tritricum aestivum, polarized light is observed in the bulbous part of the stomata reflecting radial organization around this region while the lack of polarized light in the central part of the GCs reflects the parallel orientation of the cellulose microfibril. Within sedges, stomata of (C) Cyperus esculentus stomata showing a pattern similar to dumbbell stomata, while (D) Carex flava, and (E) Schoenus nigricans, show light polarization in the central part of the stomata and at the tip/end of the bulbous part of the GCs. Radial pattern of polarized light in kidney shaped stomata as exemplified by (F) Aechmea recurvata. Scale bar in all images is equal to 10µm.

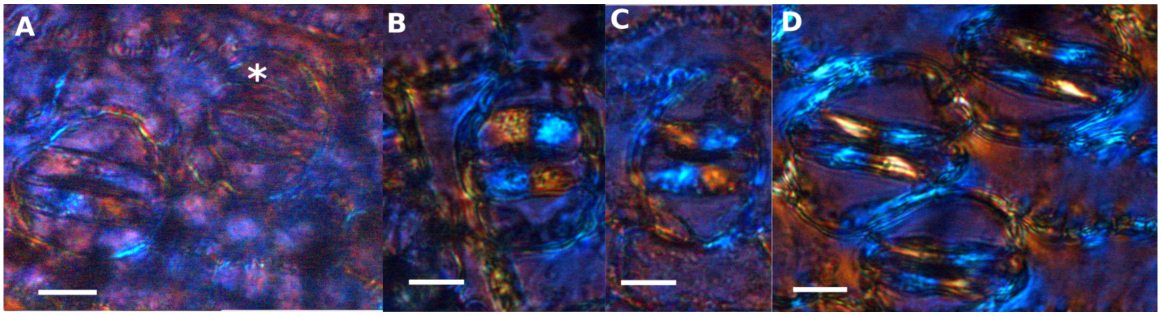


**Fig S3.** Stomatal developmental trajectory of cellulose microfibrils in *Cyperus alternifolius*. (A) Early differentiated guard cells showing low birefringence, consistent with minimal or no detectable cellulose microfibril deposition. * Denotes stomata with low birefringence (B, C) Intermediate stages of guard cell development showing cellulose microfibril organization in a kidney-like pattern. (D) Nearly mature stomata with partially rearranged cellulose microfibrils, exhibiting an intermediate organization between kidney- and dumbbell-shaped stomata. All scale bars: 20 µm.

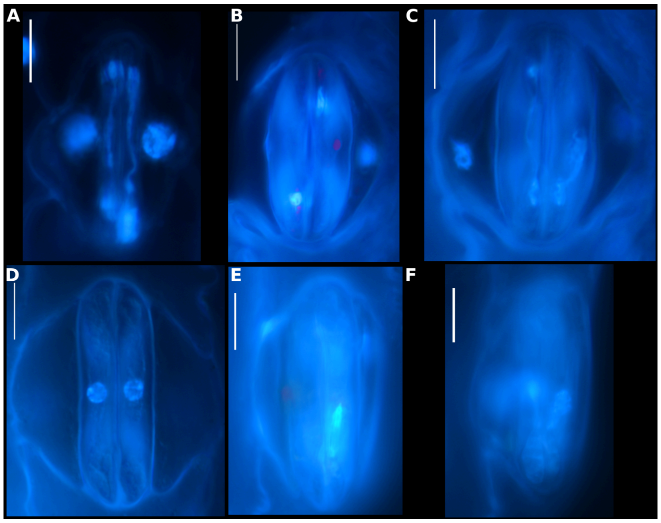


**Fig S4.** Variability in nuclear morphology in sedge species. (A) Elongated nucleus in the GCs of *Zea mays*. Note the continuity of nuclear content from one bulbous region to  the following one. (B,C) *Cyperus alternifolius* nucleus located at the bulbous regions of the GCs (B) or nucleus elongated (C) (D-E) *Cyperus esculentus* rounded nucleus located at the central region of the GCs (D) or nucleus partially elongated near the bulbous region of the GC. (F) Partially elongated nucleus located in the bulbous region of one GC of *Carex flava*. Scale bars in all images, 10 µm.

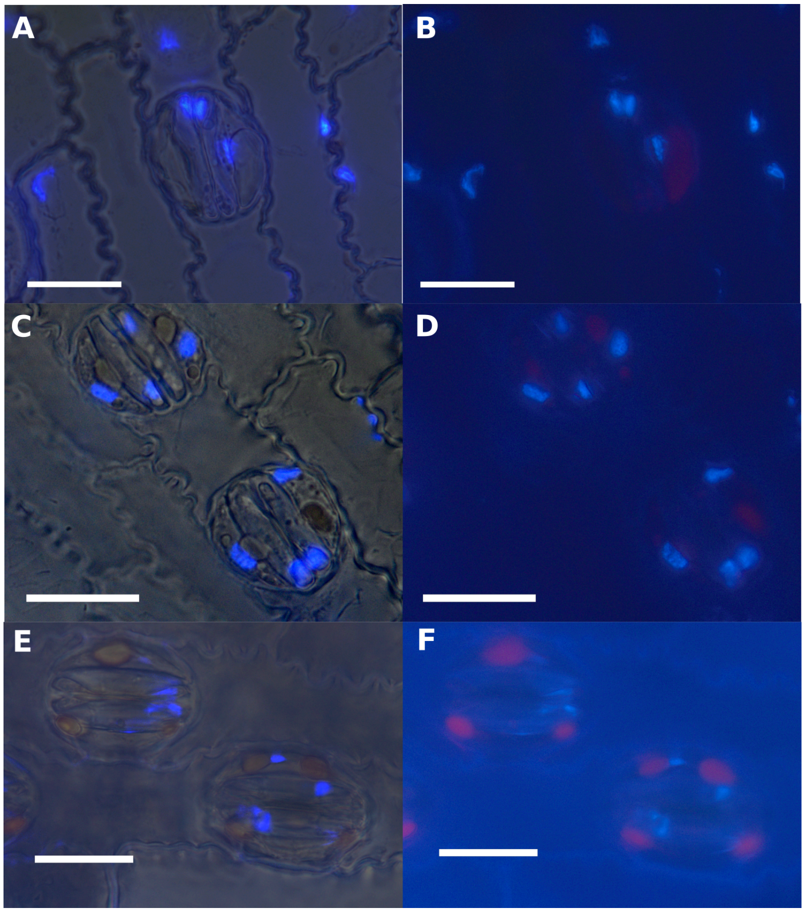


**Fig S5.** Nuclei of paired guard cells in close contact within the bulbous region of (A-D) *Carex sylvatica*, and (E,F) *C. alternifolius*. (A,C,E) Overlay of DAPI-stained fluorescence and bright-field images. (B,D,F) DAPI-stained fluorescence images. All scale bars: 20 µm.

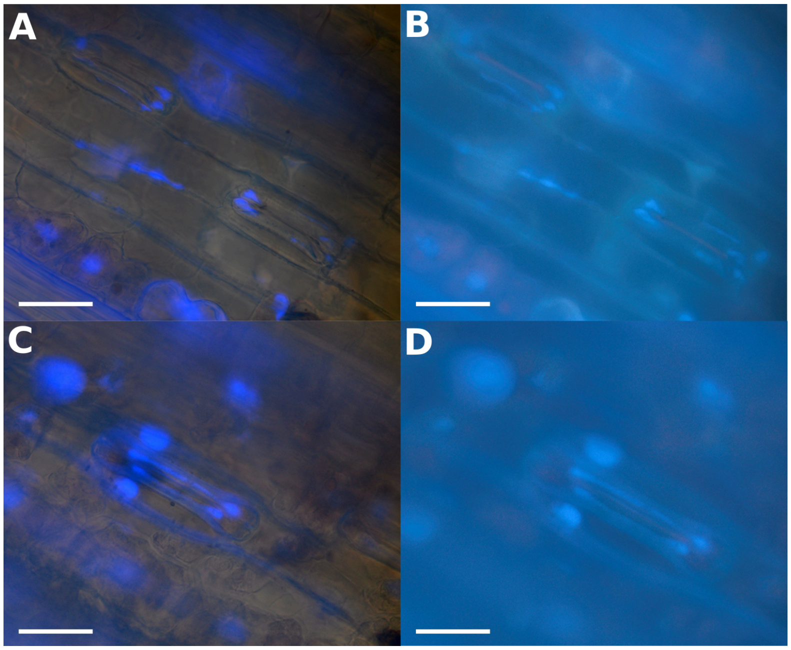


**Fig S6.** Nuclei of paired guard cells in close contact within the bulbous region of (A–D) *Lolium perenne*. (A, C) Overlay of DAPI-stained fluorescence and bright-field images. (B, D) DAPI-stained fluorescence images. All scale bars: 20 µm.

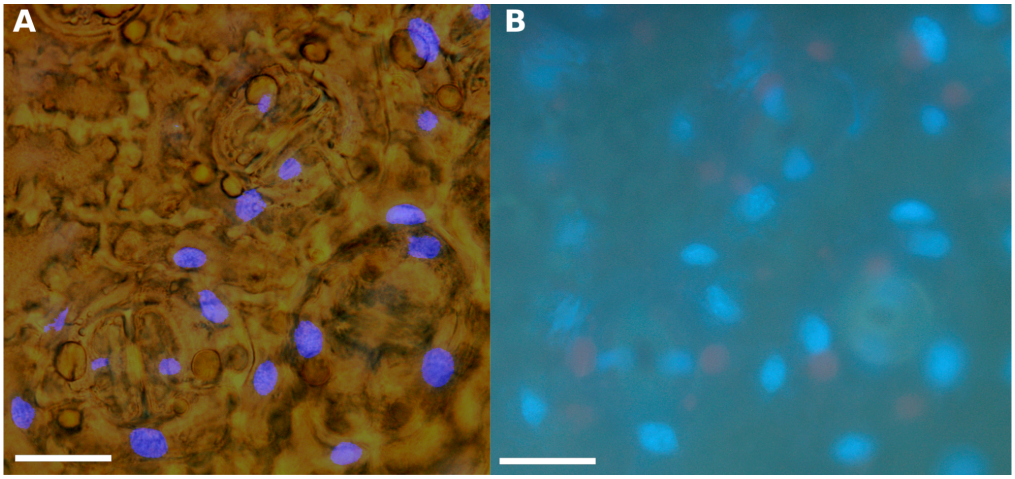


**Fig S7**. Nuclei of paired guard cells observed not in contact and far from each other in *Camellia japonica*. (A) Overlay of DAPI-stained fluorescence and bright-field images. (B) DAPI stained fluorescent image. Scale bars in all images, 20 µm.

**Methods S1**

*Image preprocessing*

To quantify the orientation of cellulose microfibrils and nuclei shape within GCs across species, we performed identical preprocessing steps. For each epidermal image, both brightfield light polarization images or brightfield and DAPI fluorescence images were opened and combined into a single stack using the *Images to Stack* function from ImageJ (Schneider et al., 2012). This allowed simultaneous operations on both channels. The region containing the target stomata was cropped, and the images were rotated to orient each stoma horizontally using the *Rotate* function. We manually outlined the GCs on the brightfield slice using the polygon selection tool, and the same regions were applied to the light polarization and the DAPI channels. The selected area was then cropped to include only the GCs, and the brightfield slice was removed, leaving a single light polarization or fluorescence image. The guard-cell selections were saved and exported as ROI files for subsequent batch processing. Fluorescence contrast was enhanced automatically (Enhance Contrast, saturation 0.35%), and the processed images were saved in TIFF format together with the corresponding ROI files.

*Stomata size measurements*

To quantify stomata size, regions of interest (ROIs) corresponding to individual stomata were converted into binary masks, in which foreground pixels represented the stomatal area. To estimate stomatal length and width, the spatial coordinates of all foreground pixels within each mask were extracted and subjected to principal component analysis (PCA). The first principal component (PC1) was taken to represent the major axis of the stomatal length, while the second principal component (PC2) represented the orthogonal minor axis. All measurements were initially obtained in pixel units and converted to micrometers depending on the pixel to micrometer relationship.

**Table S1** Taxa included in this study, describing the material source and the analyses performed for each species. Life-form descriptions follow Raunkiær's life-form classification[1]. In the Analysis column, (1) denotes cellulose microfibrils organization and (2) denotes nuclei DAPI staining.

| **Order** | **Family** | **Genus** | **Species** | **Auth.** | **Material source** | **Analysis** | **Lifeform description** | **Biome** |
|---|---|---|---|---|---|---|---|---|
| Asparagales | Amarydillaceae | *Allium* | *thunbergii* | *G.Don* | Herbarium specimen | 1 | Geophyte | Temperate |
| Poales | Bromeliaceae | *Aechmea* | *recurvata* | *(Klotzsch) L.B.Sm.* | Herbarium specimen | 1 | Epiphyte | Subtropical |
| Poales | Cyperaceae | *Carex* | *flava* | *L.* | Botanic garden of Napoli (OBN) | 1 | Hemicryptophyte or rhizomatous geophyte | Temperate |
| Poales | Cyperaceae | *Carex* | *flacca* | *Schreb.* | Seeds acquired from Jelitto Staudensamen GmbH | 2 | Hemicryptophyte or rhizomatous geophyte | Temperate |
| Poales | Cyperaceae | *Carex* | *sylvatica* | *Huds.* | Botanic garden of Zurich (Botanischer Garten UZH) | 2 | Hemicryptophyte or rhizomatous geophyte | Temperate |
| Poales | Cyperaceae | *Cyperus* | *alterniflorus* | *R.Br.* | Botanic garden of Napoli (OBN) | 1,2 | Hemicryptophyte or rhizomatous geophyte | Subtropical |
| Poales | Cyperaceae | *Cyperus* | *blepharoleptos* | *Steud.* | Herbarium specimen | 1 | Helophyte | Seasonally Dry Tropical |
| Poales | Cyperaceae | *Cyperus* | *esculentus* | *L.* | | 1,2 | Tuberous geophyte | Subtropical |
| Poales | Cyperaceae | *Cyperus* | *eragrostis* | *Lam.* | Herbarium specimen | 1 | Hemicryptophyte or rhizomatous geophyte | Seasonally Dry Tropical |
| Poales | Cyperaceae | *Cyperus* | *niveus var. leucocephalus* | *(Kunth) Fosberg* | Herbarium specimen | 1 | Hemicryptophyte or rhizomatous geophyte | Temperate |
| Poales | Cyperaceae | *Cyperus* | *pandanophyllum* | *Kunth* | Herbarium specimen | 1 | Rhizomatous geophyte | Wet Tropical |
| Poales | Cyperaceae | *Cyperus* | *papyrus* | *L.* | Botanic garden of Napoli (OBN) | 1 | Hemicryptophyte or rhizomatous geophyte | Wet Tropical |
| Poales | Cyperaceae | *Cyperus* | *schomburgkianus* | *Nees* | Herbarium specimen | 1 | Hemicryptophyte or rhizomatous geophyte | Seasonally Dry Tropical |
| Poales | Cyperaceae | *Cyperus* | *glaber** | *L.* | Seeds acquired from Jelitto Staudensamen GmbH | 2 | Hemicryptophyte or rhizomatous geophyte | Temperate |
| Poales | Cyperaceae | *Didymiandrum* | *stellatum* | *(Boeckeler) Gilly* | Palmen Garten Frankfurt | 1,2 | Hemicryptophyte or rhizomatous | Wet Tropical |

| | | | | | | | | |
|---|---|---|---|---|---|---|---|---|
| | | | | | | | geophyte | |
| Poales | Cyperaceae | *Cyperus* | *brevifolius* | *(Rottb.) Hassk.* | Herbarium specimen | 1 | Hemicryptophyte or rhizomatous geophyte | Seasonally Dry Tropical |
| Poales | Cyperaceae | *Scirpoides* | *holoschenus* | *(L.) Soják* | Botanic garden of Napoli | 1 | Hemicryptophyte | Temperate |
| Poales | Cyperaceae | *Schoenus* | *nigricans* | *L.* | Botanic garden of Napoli | 1 | Hemicryptophyte | Temperate |
| Poales | Cyperaceae | *Scirpodendron* | *ghaeri* | *(Gaertn.) Merr.* | Meise Botanic Garden | 1 | Rhizomatous geophyte or helophyte | Wet Tropical |
| Poales | Juncaceae | *Juncus* | *conglomeratus* | *L.* | Herbarium specimen | 1 | Hemicryptophyte | Temperate |
| Poales | Poaceae | *Digitaria* | *sanguinalis* | *(L.) Scop.* | Botanic garden of Napoli (OBN) | 1 | Therophyte | Temperate |
| Poales | Poaceae | *Hordeum* | *murinum* | *L.* | | 1 | Therophyte or biennial | Temperate |
| Poales | Poaceae | *Hordeum* | *vulgare* | *L.* | | 1 | Therophyte | Temperate |
| Poales | Poaceae | *Triticum* | *aestivum* | *L.* | Botanic garden of Napoli (OBN) | 1 | Therophyte or biennial | Temperate |
| Poales | Poaceae | *Zea* | *mays* | *L.* | Botanic garden of Napoli (OBN) | 1 | Therophyte | Seasonally Dry Tropical |
| Poales | Poaceae | *Lolium* | *perenne* | *L.* | | 2 | Biennial or hemicryptophyte | Temperate |
| Ericales | Theaceae | *Camellia* | *japonica* | *L.* | Botanic garden of Napoli (OBN) | 1,2 | Phanerophytes | Subtropical |
| Poales | Typhaceae | *Typha* | *laxmannii* | *Lepech.* | Herbarium specimen | 1 | Helophyte | Temperate |

**Cyperus serotinus var serotinus | Cyperus brevifolius*
